# regionReport: Interactive reports for region-based analyses

**DOI:** 10.1101/016659

**Authors:** Leonardo Collado-Torres, Andrew E. Jaffe, Jeffrey T. Leek

## Abstract

`regionReport` is a `R` package for generating detailed interactive reports from regions of the genome. The report includes quality-control checks, an overview of the results, an interactive table of the genomic regions, and reproducibility information. `regionReport` can easily be expanded with report templates for other specialized analyses. In particular, `regionReport` has an extensive report template for exploring `derfinder` results from annotation-agnostic RNA-seq differential expression analyses.

**Availability:** `regionReport` is freely available via Bioconductor at bioconductor.org/packages/release/bioc/html/regionReport.html.

## 1 Introduction

Many analyses of genomic data result in regions along the genome that associate with signal. These genomic regions can result from determining differentially bound peaks from ChIP-seq data (for example, using `DiffBind` [Stark and Brown, 2011]), identifying differentially methylated regions (DMRs) from DNA methylation data (for example, via `bumphunter` [Jaffe *et al*., 2012]), or performing base-resolution differential expression analyses using RNA sequencing data (for example, using `derfinder` [Collado-Torres *et al*., 2015]). The genomic regions themselves are commonly stored in a `GRanges` object from `GenomicRanges` [Lawrence *et al*., 2013] when working with R or the BED file format on the UCSC Genome Browser [Kent *et al*., 2002], but other information on these regions, for example summary statistics (e.g. magnitude, area, etc) and significance (e.g. p-values, false discovery rates, etc) provides useful information. The usage of R in genomics is increasingly common due to the usefulness and popularity of the Bioconductor project [Huber *et al*., 2015], and in the latest version (3.0), 181 unique packages use `GenomicRanges` for many workflows, demonstrating the widespread utility of identifying and summarizing genomic regions.

Here we introduce `regionReport` which allows users to explore genomic regions of interest through interactive stand-alone HTML reports that can be shared with collaborators. These reports are flexible enough to display plots and quality control checks within a given experiment, but can easily be expanded to include custom visualizations and conclusions. The resulting HTML report emphasizes reproducibility of analyses [Sadnve *et al*., 2013] by including all the `R` code without obstructing the resulting plots and tables. We envision `regionReport` will provide a useful tool for exploring and sharing genomic region-based results from high throughput genomics experiments.

## 2 Usage

### 2.1 Installation

`regionReport` and required dependencies can be easily installed from Bioconductor with the following commands:

~~~
source(“http://bioconductor.org/biocLite.R”)
~~~

~~~
biocLite(“regionReport”)
~~~

### 2.2 Input

The package includes a R Markdown template which is processed using `knitr` [Xie, 2013] and `rmarkdown` [RStudio Inc., 2014], then styled using `knitrBootstrap` [Hester, 2014].

To generate the report, the user first has to identify the regions of interest according to their analysis workflow. For example, by performing bumphunting to identify DMRs with `bumphunter`. The report is then created using `renderReport()` which is the main function in this package. The argument `customCode` can be used to customize the report if necessary.

For the `derfinder` use case, the `derfinderReport()` function creates the recommended report that includes visualizations of the coverage information for the best regions and clusters of regions.

### 2.3 Output

A small example can be generated using:

~~~
example(“renderReport”, “regionReport”, ask=FALSE)
~~~

The Supplementary Website contains reports using `BiffBind, bumphunter` and `derfinder` results. The `derfinder` use case is illustrated with data sets previously described [Collado-Torres *et al*., 2015] which span simulation results, a moderately sized data set (25 samples), and a large data set with 487 samples; thus covering a wide range of scenarios.

Note that alternative output formats such as PDF files are also available, although they are not as dynamic and interactive as the HTML format.

## 3 Report

The report includes a series of plots for checking the quality of the results and browsing the table of regions. Each element has a brief explanation, although actual interpretation of the results is data set and workflow-dependent. The code for each plot or table is hidden by default and can be shown by clicking on the appropriate toggle. To facilitate navigation a menu is always included, which is useful for users interested in a particular section of the report.

### 3.1 Quality checks

This section of the report includes a variety of quality control steps which help the user determine whether the results are sensible. The quality control steps explore:

- P-values, Q-values, and FWER adjusted p-values
- Region width
- Region area: sum of single-base level statistics (if available)
- Mean coverage or other score variables (if available)

A combination of density plots and numerical summaries are used in these quality checks. If there are statistically significant regions, the distributions are compared between all regions and the significant ones. For example, the distribution region widths might have a high density of small values for the global results, but shifted towards higher values for the subset of significant regions.

### 3.2 Genomic overview

The report includes plots to visualize the location of all the regions as well as the significant ones. Differences between them can reveal location biases. The nearest known annotation feature for each region is summarized and visually inspected in the report.

### 3.3 Best regions

An interactive table with the top 500 (default) regions is included in this section. This allows the user to sort the region information according to their preferred ranking option. For example, lowest p-value, longest width, chromosome, nearest annotation feature, etc. The table also allows the user to search and subset it interactively. A common use case is when the user wants to check if any of the regions are near a known gene of their interest.

### 3.4 Reproducibility

At the end of the report, detailed information is provided on how the analysis was performed. This includes the actual function call to generate the report, the path where the report was generated, time spent, and the detailed R session information including package versions of all the dependencies.

The `R` code for generating the plots and tables in the report is included in the report itself, thus allowing users to manually reproduce any section of the report, customize them, or simply change the graphical parameters to their liking.

### 3.5 derfinder use case: coverage plots

In the `derfinder` use case, for each of the best 100 (default) DERs a plot showing the coverage per sample is included in the report. These plots allow the user to visualize the differences identified by `derfinder` along known exons, introns and isoforms. The plots are created using `derfinderPlot`, also available via Bioconductor.

Due to the intrinsic variability in RNA-seq coverage data or mapping artifacts, two candidate DERs may be called by `derfinder` that are relatively close and there might be reasons to consider them as a single candidate DER. This tailored report groups candidate DERs into clusters based on a distance cutoff. After ranking them by their area, for the top 20 (default) clusters it plots tracks with the coverage by sample, the mean coverage by group, the identified candidate DERs colored by whether they are statistically significant, and known alternative transcripts.

## 4 Conclusions

`regionReport` creates interactive reports from a set of regions and can be used in a wide range of genomic analyses. Reports generated with `regionReport` can easily be extended to include further quality checks and interpretation of the results specific to the data set under study. These shareable documents are very powerful when exploring different parameters values of an analysis workflow or applying the same method to a wide variety of data sets. The reports allow users to visually check the quality of the results, explore the properties of the genomic regions under study, inspect the best regions and interactively explore them.

Furthermore, `regionReport` promotes reproducibility of data exploration and analysis. Each report provides R code that can be used as the starting point for other analyses within a dataset. `regionReport` provides a flexible output for exploring and sharing results from high throughput genomics experiments.

~~~
*Suplementary Website* http://lcolladotor.github.io/regionReportSupp/
~~~

*Funding* J.T.L. was supported by NIH Grant 1R01GM105705, L.C.T. was supported by Consejo Nacional de Ciencia y Tecnología México 351535.

## References

Collado-Torres, L. et al. (2015) derfinder: Software for annotation-agnostic RNA-seq differential expression analysis, bioRxiv, 015370.

Jaffe, AE et al. (2012) Bump hunting to identify differentially methylated regions in epigenetic epidemiology studies, International journal of epidemiology, 41(1), 200–209.

Hester, J. (2014), knitrBootstrap: Knitr Bootstrap framework, R package version 1.0.0, github.com/jimhester/knitrBootstrap.

Huber, W et al. (2015), Orchestrating high-throughput genomic analysis with Bioconductor, Nature Methods, 12, 115–121.

Kent WJ et al. (2002) The human genome browser at UCSC, Genome research, Jun;12(6):996–1006.

Lawrence, M. et al. (2013), Software for computing and annotating genomic ranges, PLoS computational biology.

RStudio Inc. (2014), rmarkdown: Dynamic Documents for R, R package version 0.1.90, rmarkdown.rstudio.com.

Sandve, GK. et al. (2013), Ten simple rules for reproducible computational research, PLoS computational biology.

Stark R and Brown G (2011). DiffBind: differential binding analysis of ChIP-Seq peak data, R package version 1.12.3, bioconductor.org/packages/release/bioc/vignettes/DiffBind/inst/doc/DiffBind.pdf.

Xie, Y. (2013), Dynamic Documents with R and knitr. Chapman and Hall/CRC, Boca Raton, Florida. ISBN 978-1482203530.

